# *ALI-1*, candidate gene of *B1* locus, is associated with awn length and grain weight in common wheat

**DOI:** 10.1101/688085

**Authors:** Dongzhi Wang, Kang Yu, Di Jin, Linhe Sun, Jinfang Chu, Wenying Wu, Peiyong Xin, Xin Li, Jiazhu Sun, Wenlong Yang, Kehui Zhan, Aimin Zhang, Dongcheng Liu

## Abstract

Awn plays a vital role in the photosynthesis, grain production and drought tolerance of common wheat; however, works on the systematic identification or cloning of genes controlling wheat awn length (AL) were seldom reported. Here, we conducted the Genome-wide association study (GWAS) in 364 wheat accessions and identified 25 loci involved in the AL, including dominant awn suppressors *B1*, *B2* and four homologs of awn controlling genes in rice and barley. Furthermore, the *B1* locus was mapped to a 125-kb physical interval harboring two genes on chromosome 5AL through map-based cloning. As the candidate gene for *B1* locus, a C_2_H_2_ zinc finger gene *Awn Length Inhibitor 1* (*ALI-1*) expressed predominantly in the developing spike of awnless individuals and suppresses downstream genes transcriptionally. *ALI-1* reduces cytokinin content and simultaneously restrains cytokinin signal transduction, which leads to a stagnation of cell proliferation and reduction of cell number in awn. Noteworthily, *ali-1* was the first awn controlling locus that observed increasing grain length in wheat, which is a valuable supplemental attribution of awn on grain weight besides photosynthesis. Thus, *ALI-1* pleiotropically regulates awn and grain development, and this work provides a strategy to achieve improved grain yield and address future extreme climate.

**Highlight:** *ALI-1*, candidate gene of awn suppressing *B1* locus, associates with awn length and grain length, providing a reacquaint of the effect of wheat awn on grain production.

## Introduction

As an important component of the spike, awn is a long needle-like apical extension of the lemma formed on the florets of grass species, e.g., wheat, barley, rye, oats, sorghum and rice. Wheat awns have an acutely triangular shape with sclerenchyma, two chlorenchyma zones and three well-developed vascular bundles inside, and rows of stomata on the epidermis of the abaxial terminal (Li et al., 2010). The photosynthesis of chlorenchyma in awn serves as an essential supplemental assimilates for grain filling, especially when flag leaves senesce (Grundbacher, 1963; Olugbemi et al., 1976; Weyhrich et al., 1994; Li et al., 2006). Under drought condition, awn contributes up to 16% of the total grain weight and production (Thorne, 1965; Evans et al., 1972; Duwayri, 1984; Blum, 1985; Maydup et al., 2010). Besides, long awn in the wild wheat and its relatives protects seeds from shattering and predation, facilitates seed dispersal, helps balance and land the embryo, and propels seed burial (Grundbacher, 1963; Sorensen, 1986; Elbaum et al., 2007; Hua et al., 2015).

Many genes involved in the development of awn have been cloned in cereal crops, such as *Lks2*, *Hooded* and *ROUGH AWN1* in barley, and *An-1*, *An-2/LABA1*, *GAD1/RAE2*, *DL*, *GLA*, *OsETT2* and *SHL2* in rice (Müller et al., 1995; Yuo et al., 2012; Luo et al., 2013; Toriba and Hirano, 2014; Hua et al., 2015; Gu et al., 2015; Jin et al., 2016; Bessho-Uehara et al., 2016; Milner et al., 2018; Zhang et al., 2019). In common wheat, the elaborately genome-wide identification of genes controlling the awn length was seldom reported, although there are a few works on the genome-wide association analysis of awn presence/absence or awn type (Sheoran *et al.*, 2019; Mackay et al., 2014; Liu et al., 2017a). *Tipped1* (*B1*), *Tipped2* (*B2*) and Hooded (*Hd*) are the known dominant genes suppressing awn development in common wheat and whose different combinations bring in variations for awn performance (Watkins and Ellerton, 1940). The *B1* produces apically tip-awned phenotype with short awns at the top and absent at the base and middle of the spike (Watkins and Ellerton, 1940). The awn tips of *B1* are usually straight and unbent at the base, while awn in *B2* is gently curved and nearly equal in length along the spike. For the *Hd*, awns are reduced in length, curved/twisted and in some cases considerably broadened at the base resembling a membranous lateral expansion of *Hooded* mutants in barley (Watkins and Ellerton, 1940; Müller et al., 1995). *B1* was located on the long arm of chromosome 5A and narrowed to a 7.5-cM interval closely linked with marker BW8226_227 (Sourdille et al., 2002; Mackay et al., 2014; Yoshioka et al., 2017), while *B2* and *Hd* were on the long arm of 6B and the short arm of 4A, respectively (Sourdille et al., 2002; Yoshioka et al., 2017). However, none of the causative genes have been cloned, fine mapped, or molecularly systematic investigated.

Given the potential influence of awns on yield potential and drought tolerance, breeding wheat varieties with long awns especially in Europe where varieties are predominantly awnless, might help to deal with the threat of future food crisis and climate change. A better understanding of the genetics, evolution, and molecular mechanism of awn would facilitate this process. Here, a whole-genome-wide identification of genes controlling the awn length in common wheat was performed, and a candidate gene conferring to the awn suppressing of *B1* locus was characterized.

## Materials and methods

### Plant materials

Accessions in GWAS panel (**Table S1**) were grown in Zhaoxian (37°51′N, 114°49′E) during three successive cropping seasons (2015-2018) in individual (R1) and plots (R2). The field experiment was performed using a completely randomized design, and each accession planted in six 200-cm-long rows. Agronomic management followed local practices.

Several F_12:13_ populations derived from the NongKeYuanSanLiMai/NongDa3214 cross were identified using tightly linked SSR markers and served as a set of near-isogenic lines (NILs) of *ALI-1.* The NILs were planted in Zhaoxian during the 2016–2017 and 2017–2018 cropping seasons with regular management and drought treatment, respectively.

### Phenotypic evaluation

Six awns at the middle of spikes each for five spikes were averaged, representing the awn length of each accession. Agronomical traits were measured accordingly, and kernel traits investigated using the SC-G image analysis system (Wanshen Detection Technology Co., Ltd., Hangzhou, China).

### Genome Wide Association Analyses

Each accession was genotyped using Affymetrix Wheat660K SNP arrays by Capital Bio Corporation (Beijing, China). GWAS was performed using Tassel v5.2 with single chromosomal-located SNPs on IWGSC RefSeq V1.0 after quality control (missing rate ≤ 10% and MAF ≥ 5%). Significant markers were visualized by Manhattan plots and quantile-quantile plots using the R package “qqman” A significance threshold of –log_10_*P* ≥ 3.5 was applied to declare significant SNPs. The pairwise *r^2^* (squared allele frequency correlation) values were calculated and displayed with LD plots by Haploview 4.2 software (Barrett et al., 2005).

### Primers

The primers used in this study are listed in **Table S10**.

### Molecular mapping

SSR markers were designed in SSRLocator, and CAPS and dCAPS primers in the CAPS/dCAPS Designer, respectively, and the resultant genotypes were subjected to genetic linkage map construction in JoinMap 4 (Van Ooijen J. 2006.) and drawn using Mapchart v2.3 software (Voorrips, 2002).

### Sequencing and Data Analysis

Genomic DNA region of *TraesCS5A02G542800* and *TraesCS5A02G542900* in each accession were amplified using primer pairs (**Table S10**), with 16-nt asymmetric barcodes tagged on the 5′ end (https://github.com/PacificBiosciences/Bioinformatics-Training/wiki/Barcoding-with-SMRT-Analysis-2.3). The library of purified PCR products pool was sequenced using a PacBio RS II SMRT DNA Sequencing System (Kozich et al., 2013). The output circular consensus sequencing reads were assigned to each accession separated by barcode sequences, and aligned to the gene reference sequence in Chinese Spring using the software BWA. Randomly selected sequence variations were verified by realigned using Clustal-W and confirmed using Sanger sequencing in some accessions.

### Phylogenetic analysis

Protein sequences of *TraesCS5A02G542800* and all C_2_H_2_ genes in *Arabidopsis thaliana* were performed to the phylogenetic reconstruction in MEGA version 7.0.26 using the neighbor-joining method. Bootstrap values were estimated (with 1000 replicates) to assess the relative support for each branch.

### RNA-Seq analyses

Young spikes (1.0-2.0 cm in length) of homozygous dominant and recessive individuals (genotyped by SSR88, SSR151 and InDel-07) from three pairs of NILs were harvested and pooled into six samples (three lines × two genotypes, ≥ 200 spikes per sample). Each sample was evenly divided into two portions for RNA isolation and quantitative content determination of endogenous CKs and IAA.

Total RNA was extracted using a TRIzol kit (Invitrogen) and sequenced by the BGI (Shenzhen, China) on HiSeq 4000 (Illumina, San Diego, USA). Filtered reads were mapped to Chinese Spring TGAC v1 genome assembly (http://plants.ensembl.org/Triticum_aestivum), and transcripts aligned to each gene was calculated and normalized to FPKM values. Significant differentially expressed genes (DEGs) in each pair of NILs were screened through NOISeq, with a threshold of |log2(FPKM_Awnless_/FPKM_Awned_)| ≥ 1 and probability ≥ 0.8 (Tarazona et al., 2011). The enrichment of GO terms and the Kyoto Encyclopedia of Genes and Genomes (KEGG) pathway were conducted using R package clusterProfiler (Yu et al., 2012).

### Real-time PCR analysis

Quantitative real-time PCR was performed on a LightCycler 480 system (Roche, Indianapolis, IN, USA) using *Ta4045* gene as internal reference (Paolacci et al., 2009). The comparative CT method (ΔΔCT) was used in the quantification analysis (Benjamini and Hochberg, 1995).

### Quantitative analyses of endogenous CKs and IAA

Quantitative analyses of endogenous CKs and IAA were conducted based on the method reported previously, with three biological replicates (Du et al., 2017; Fu et al., 2012). The ^2^H_2_-IAA was served as the internal standard of IAA, while D_5_-*t*Z, D_5_-*t*ZR, D_6_-iP and D_6_-iPR were used as the internal standards for CKs. LC-MS/MS analysis was performed with purified extracts on an ACQUITY UPLC system coupled to the 6500 Q-Trap system (AB SCIEX).

### Transcriptional activation analysis of *ALI-1*

The GAL4 reporter plasmid was generated using the firefly LUC reporter gene driven by the minimal TATA box of the 35S promoter plus five GAL4 binding elements, and the ORF of *ALI-1* amplified by PCR were fused into the Pro*_35S_*:GAL4DBD vector to construct the effecter plasmid. The Pro*_35S_*:GAL4DBD vector was used as negative control and the Pro*_35S_*:GAL4DBD:VP16 vector fused with strong activation protein VP16 as a positive control. Other procedures carried out following the protocol reported previously with six independent measures carried out for each analysis (Hao et al., 2010).

### Histological observations

Awns with lemma of awned and awnless individuals (5 ± 0.5 cm in spike length) were fixed with FAA solution, embedded in paraffin, then longitudinally sectioned, stained with 1.0% Safranin O and 0.5% FastGreen, and observed using NIKON CI-S microscope. The cell lengths of each sample were measured on three serial sections at the upper, middle and bottom parts of awn. Cell number for the entire length of an awn was estimated based on the length of awns.

### Statistical analysis

Descriptive statistics, analyses of variance (ANOVA), Pearson’s correlation analyses were performed using OriginPro, Version 2019 (OriginLab Corporation, Northampton, MA, USA). Variance components were used to calculate broad sense heritability (*h^2^*) of awn length defining as *h^2^*=σ*_g_^2^*/(σ*_g_^2^*+σ*_ge_^2^*/*r*+σ_ε_*^2^*/*re*) (Liu et al., 2017a). To eliminate the environmental impact, the BLUP value across all tested environments was calculated using R package “lme4”.

## Results

### 25 loci including *B1* significantly associated with wheat awn length in the Genome-wide-association panel

During the 2015–2016, 2016–2017 and 2017–2018 cropping seasons, winter wheat (*Triticum aestivum* L.) panel of 364 accessions were grown at Zhaoxian in individual (R1) and plots (R2) (**Table S1**). Significant variation (*P* < 0.001) of AL was identified among the 364 accessions across all six environments, ranging from 0 mm to 110 mm with a coefficient of variation ranged from 0.22 to 0.26 (**Table S2**, **Figure S1**), indicating that this population embodies abundant variations and suitable for the GWAS. A significant correlation was detected between environments with Pearson’s correlation coefficients between 0.61–0.91 (**Figure S1**). Significant differences (P ≤ 0.001) among genotypes, environments, and genotype × environment interactions were observed with ANOVA (**Table S3**), and a high broad sense heritability (*h^2^*=σ*_g_^2^*/(σ*_g_^2^*+σ*_ge_^2^*/*r*+σ_ε_*^2^*/*re*), *h^2^=*0.985) was observed across all the six environments, revealing that major phenotypic variation was derived from genetic factors.

A total of 439,209 SNPs from Affymetrix 660K SNP arrays were subjected to the GWAS analysis on AL using the mixed linear model. SNP cluster (more than three SNPs with −log_10_(*P*-value) ≥ 3.50 in less than 1 Mb distance) detected in resulting best linear unbiased predictors value (BLUP) and at least three environments was regarded as a reliable significant association locus (SAL). This allowed 25 SAL associated with AL on chromosome 1A, 1D, 2A (2), 2B (3), 3A (2), 3B (2), 3D, 4A, 4B, 5A (3), 5B (2), 6B (2), 7A (2) and 7D (2), explaining phenotypic variation of BLUP ranging from 5.91% to 13.99%, respectively (**Figure 1a,b and Figure S2**, **Table S4**). These SAL were compared with previously reported genes, QTL, or markers of awn controlling loci based on the physical positions on IWGSC RefSeq V1.0 of Chinese Spring (IWGSC, 2018). Four SAL, *AX-95086847*, *AX-111043485*, *AX-109508056* and *AX-108780287* were overlapped with the homologs of *An-1*, *OsETT2*, *SHL2* in rice and *Lks2* in barley, respectively. *AX-109312058* and *AX-109882617* were with wheat awn inhibiting loci *B1* and *B2*, respectively.

**Figure 1.**
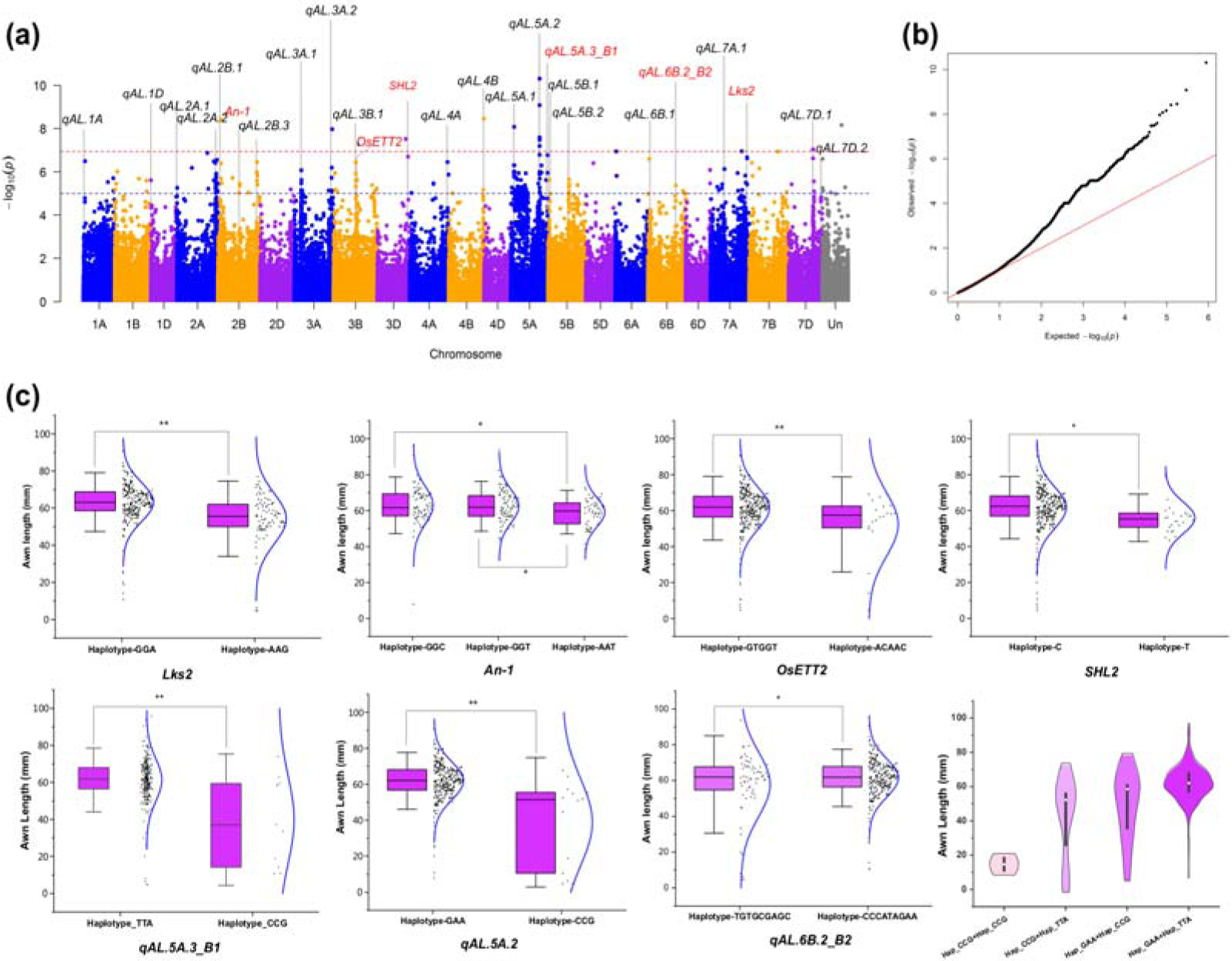
Genome-wide association study of awn length among the 364 wheat accessions. (a) Manhattan plots for BULP value of AL identifies 25 SAL across the 21 chromosomes using the mixed linear model (MLM). The −log_10_(*P*) values from a genome-wide scan are plotted against positions on 21 chromosomes. Blue and red horizontal dashed lines indicate the genome-wide significance threshold of *P*=10^−5^ and *P*=10^−7^, respectively. Gray solid vertical lines were used to depict the QTL of AL. Labels in red and black indicate the QTL overlapping with reported gene/QTL and new QTL, respectively. (b) Quantile-quantile plot of MLM for AL. The solid red line indicates the expected values. (c) Haplotypes and their distribute frequency of SAL among the wheat natural population. The boxes cover the twenty-fifth to seventy-fifth percentiles with a middle line indicates median, the whiskers outside the box extend to the ±1.5 SD. AL of each accession and their normal distribution are displayed by the box using black dots and blue curve. The differences of mean values among haplotypes were tested using the Fisher LSD test. *, *P* < 0.05, **, *P* < 0.01. The violin plot shows a smoothed approximation of the frequency distribution (a kernel density plot) was used to compare haplotype combinations of *qAL.5A.2* and *qAL.5A.3_B1*, a standard box plot is represented within the violin plot, with the mean value of the distribution shown as a white dot. The Tukey’s original box plot and the violin plot were plotted using software OriginPro, Version 2019.

Haplotypes of the significant SNPs among the GWAS accessions were identified based on the genotypes of these SAL, and the effects on AL for each haplotype were calculated (**Figure 1a**, **Table S5**). For the *Lks2*, haplotype-AAG included 91 accessions with an average AL of 54.28 mm, reducing AL for 8.97 mm as comparing with haplotype-GGA (267 accessions, 63.25 ± 10.58 mm) (**Figure 1c**, **Table S5)**. Similarly, elite haplotypes (with shorter awn) of *OsETT2*, *An-1, SHL2, qAL.5A.3_B1*, and *qAL.6B.1_B2* reduced AL by 8.99, 3.58, 5.62, 21.46 and 3.61 mm, respectively (**Figure 1c**, **Table S5**). Beyond that, the other 18 SAL has not been reported and might be potential loci controlling AL in common wheat. Among these SAL, the *qAL.5A.2* explained the most phenotypic variation (BLUP, 13.99%) and reduced AL by 23.02 mm (**Figure 1c**, **Table S5**). Interestingly, 18 out of 46 genes in the *qAL.5A.2* LD block (547.59–548.25 Mb) are auxin-responsive proteins, suggesting an auxin-mediated potential mechanism of this locus on AL **(Table S6**). Among all SAL, *qAL.5A.2* and *qAL.5A.3_B1* explain 43.15% phenotypic variation, reducing the AL from 62.09 mm (Hap_GAA+Hap_TTA, double inferior haplotype) to 14.63 mm (Hap_CCG+Hap_CCG, double elite haplotype) (**Figure 1c**). Since *B1* was a major and well-known awn controlling locus in common wheat, fine mapping and candidate gene characterization of *qAL.5A.3_B1* was further carried out.

### The *qAL.5A.3_B1* was fine mapped to a 125 kb region

SNPs in the 2 Mb regions around AX-109312058, the representative SNP of *qAL.5A.3_B1* **(Table S7**), were used to calculate the pairwise *r^2^* values. LD plot formed a ∼0.14 Mb LD block (*AX-86177799*–*AX-109312058*, 698.00–698.14 Mb) and a ∼0.94 Mb one (*AX-110564755*–*AX-109843442*, 698.18–699.12 Mb), illustrating that the *qAL.5A.3_B1* was mapped to the 698.00–699.12 Mb interval (**Figure S3**).

Based on the awn performance and genotype at *qAL.5A.3_B1* locus, two bi-parental genetic populations YS-F_2_ and NN-F_2_ derived from the crosses of YeMaiZi (YMZ, awnless) with Shi4185 (S4185, awned) and NongKeYuanSanLiMai (NK, awnless) with NongDa3214 (ND3214, awned) respectively were developed to fine map the *B1* locus (**Figure 2a,b**). The AL of F_1_ resembled the awnless parents and individuals in the F_2_ populations were divided into awnless or awned groups (**Figure 2b**), the segregation of awnless to awned individuals fit the expected ratio of 3:1 in both F_2_ populations (YS, χ^2^ = 0.44, *P* = 0.50; NN, χ^2^ = 0.88, *P* = 0.35), demonstrating that the performance of the awn inhibition was controlled by a single dominant gene. The segregation in subsequently derived YS-F_2:3_ and YS-F_3:4_ also agreed with expected Mendelian inheritance ratios of 3:1 (**Table S8**).

**Figure 2.**
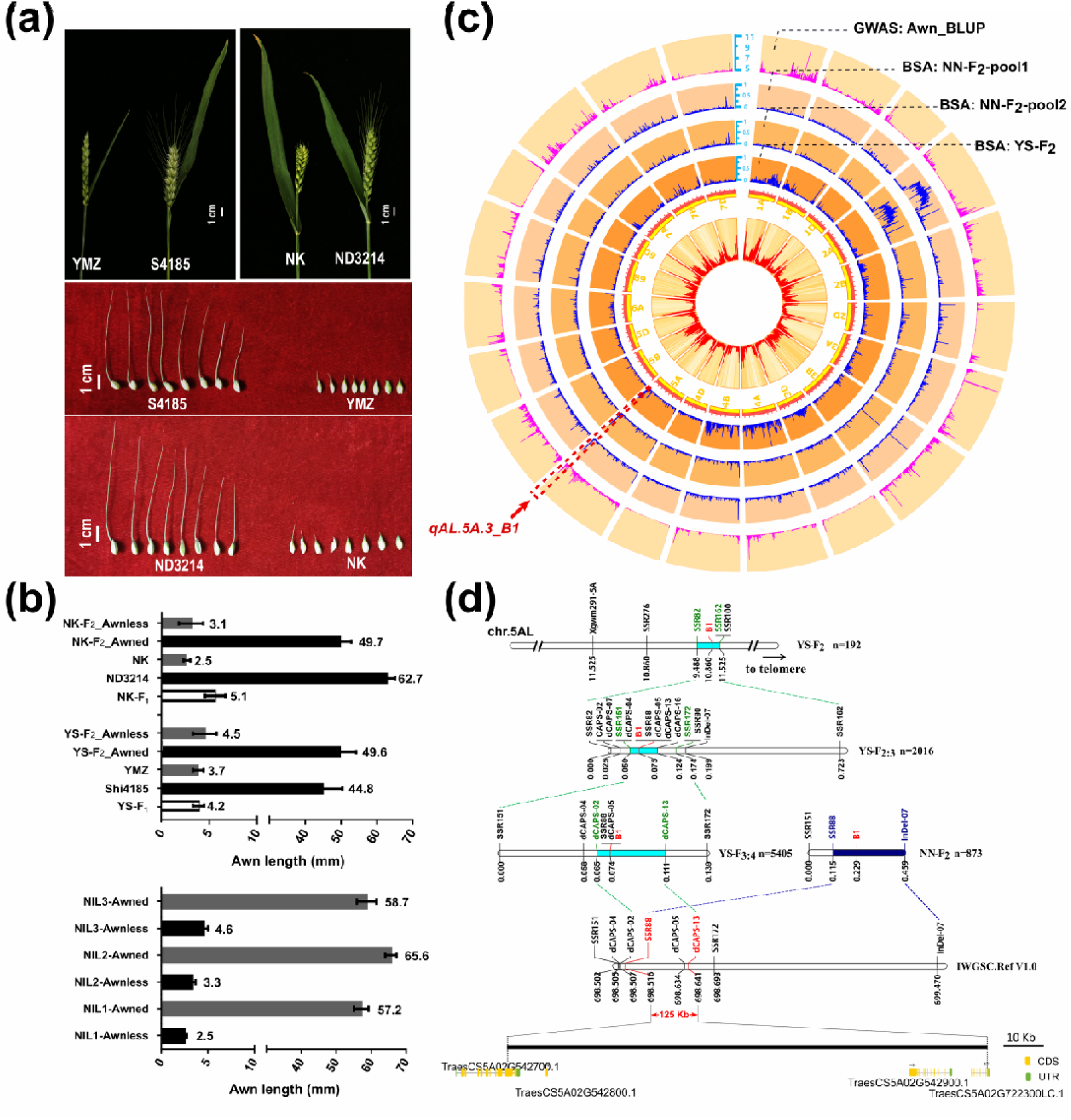
The *qAL.5A.3_B1* locus was fine mapped to a 125 kb interval harboring two genes. (a,b) The awn performance of YS-F_2_, NN-F_2,_ and NILs. Spikes of parents YMZ/S4185 (YS-F_2_ population) and NK/ND3214 (NN-F_2_ population and NILs) were displayed (a) and the AL in the YS-F_2_ population, NN-F_2_ population and NILs were measured (b). (c) A circos plot indicating BSA enrichment peaks overlapping with *qAL.5A.3_B1* locus. The relative frequency distribution of chip SNPs per 1 Mb in each chromosome was displayed in the innermost cycle using heatmap and histogram plot. The density of SNPs positively correlated with the color depth in the heatmap. The second inner circle indicates 21 chromosomes of wheat, with the physical location marked in a scale of 10 Mb. The histogram plot of associated SNPs in the GWAS (using −log_10_(*P*-values) of SNP in BLUP data) was displayed in the outermost layer, and the histogram plots indicate the distribution of polymorphic SNPs frequency on each chromosome in NN-F_2_-Pool1, NN-F_2_-Pool2 and YS-F_2_were displayed in outer layer 2–4, respectively. The polymorphic SNPs frequency was defined as the ratio of polymorphic SNPs in the total SNPs per 1 Mb in each chromosome. The *qLA.5A.3_B1* region was surrounded by a red dotted box and indicated by an arrow. The circos plot was drawn in TBtools. (d) The fine mapping of *B1* locus using bi-parental mapping populations. The physical locations of markers were marked on the physical map on chromosome 5A of Chinese Spring IWGSC RefSeq V1.0. The linkage map of each population and the physical map were lined together by consensus genetic markers using dashed lines, and the *B1* mapping intervals in each population were filled with cyan or blue. The genes in and around the 125 kb *B1* interval were displayed.

To fine map the *B1* locus, *Xgwm291-5A*, previously reported linkage marker of *B1* were subjected to screen 12 long-awn and 12 awnless individuals (Kosuge et al., 2008). The data provided that the *B1* locus should be the causative factor for the awn presence/absence in both YS-F_2_ and NN-F_2_ population. Bulk separating analysis of wheat660K SNP chip with awned and awnless pools in YS-F_2_ and NN-F_2_ population also detected a significantly differential marker enrichment at the distal end of chromosome 5AL, overlapping with the *qAL.5A.3_B1* locus (**Figure 2c**). Three 1-Mb genome sequences with 8 Mb in-between on IWGSC RefSeq V1.0 Chromosome 5A were subjected to SSR marker development, and resultant polymorphic markers *SSR276*, *SSR82*, *SSR162*, and *SSR100* were screened using the whole YS-F_2_ population. With this, *B1* was mapped to the *SSR82*–*SSR162* interval (**Figure 2d**). The derived YS-F_2:3_ population was screened with flanking marker *SSR82* and *SSR162*, and the resultant 23 recombinants between *SSR82* and *SSR162* were subjected to further analysis. More SSR markers and InDel markers based on gene sequencing were developed in the *SSR82*–*SSR162* interval, and SNPs between bulks in YS-F_2:3_ population were performed to design CAPS/dCAPS primers. With this effort, the *B1* region was narrowed to 0.074 cM interval flanked by *SSR151* and *dCAPS-06* (**Figure 2d**). With a larger YS-F_3:4_ population composed of 5405 individuals, *B1* was flanked by proximal marker *dCAPS-02* and distal marker *dCAPS-13* and co-segregated with *SSR88* and *dCAPS-05.* Thus, the *B1* region was narrowed to a 0.046 cM interval, corresponding to a 134 kb physical region on IWGSC RefSeq V1.0 chromosome 5A (**Figure 2d**).

To confirm the mapping region in YS-F_3:4_ population, the NN-F_2_ population were screened with *SSR151*, *SSR88* and *InDel-07*. One recombination event between *SSR88* and *InDel-07* was identified, and this allows the *B1* locus delimited to a 125 kb interval (698.516–698.641 Mb) (**Figure 2d**). According to the IWGSC RefSeq V1.1 annotation (https://wheat-urgi.versailles.inra.fr/Seq-Repository/Annotations), this interval only harbors two genes, *TraesCS5A02G542800* and *TraesCS5A02G542900* (**Figure 2d**), one of which should be the candidate gene of the *B1* locus.

### A C_2_H_2_ zinc finger *ALI-1* is the candidate gene for the *B1* locus

To assess the gene expression patterns of candidate genes for the *B1* locus, the expression databases of “Developmental time-course of Chinese Spring” and “Developmental time-course of Azhurnaya” through Wheat Expression Browser (http://www.wheat-expression.com/) were searched (Borrill et al., 2016; Ramírez-González et al., 2018). The homolog genes of *TraesCS5A02G542800* are dynamically expressed at different stages and tissues, predominantly in the spike, while *TraesCS5A02G542900* does not have any homolog expression bias and apparent tissue specificity (**Figure 3a, b**). Besides, analysis of the transcriptome profiling during spike development in KN9204 demonstrated that *TraesCS5A02G542800* is expressed in the early stages of spike development before the glume primordium differentiation stage (Li et al., 2018), from which period the formation of lemma starts and difference between awned and awnless individuals emerges (Luo et al., 2013; Vahamidis et al., 2014), but expression of *TraesCS5A02G542900* basically remains unchanged throughout the spike development process (**Figure 3c**). Through quantitative real-time PCR analysis with the developing spikes of three *B1* NILs, *TraesCS5A02G542800* were significantly up-regulated (23.28, 5.63 and 5.28 folds) in awnless lines, while the expression of *TraesCS5A02G542900* could not provide consistent data among these NILs (**Figure 3d**). Moreover, RNA-Seq analysis of developing spikes revealed that *TraesCS5A02G542800* had a much higher expression level than its homolog genes, *TraesCS4B02G345000* and *TraesCS4D02G340000*. More importantly, its expression in awnless lines was up-regulated with 4.3-8.7 folds as compared to the awned lines (**Figure 3e**).

**Figure 3.**
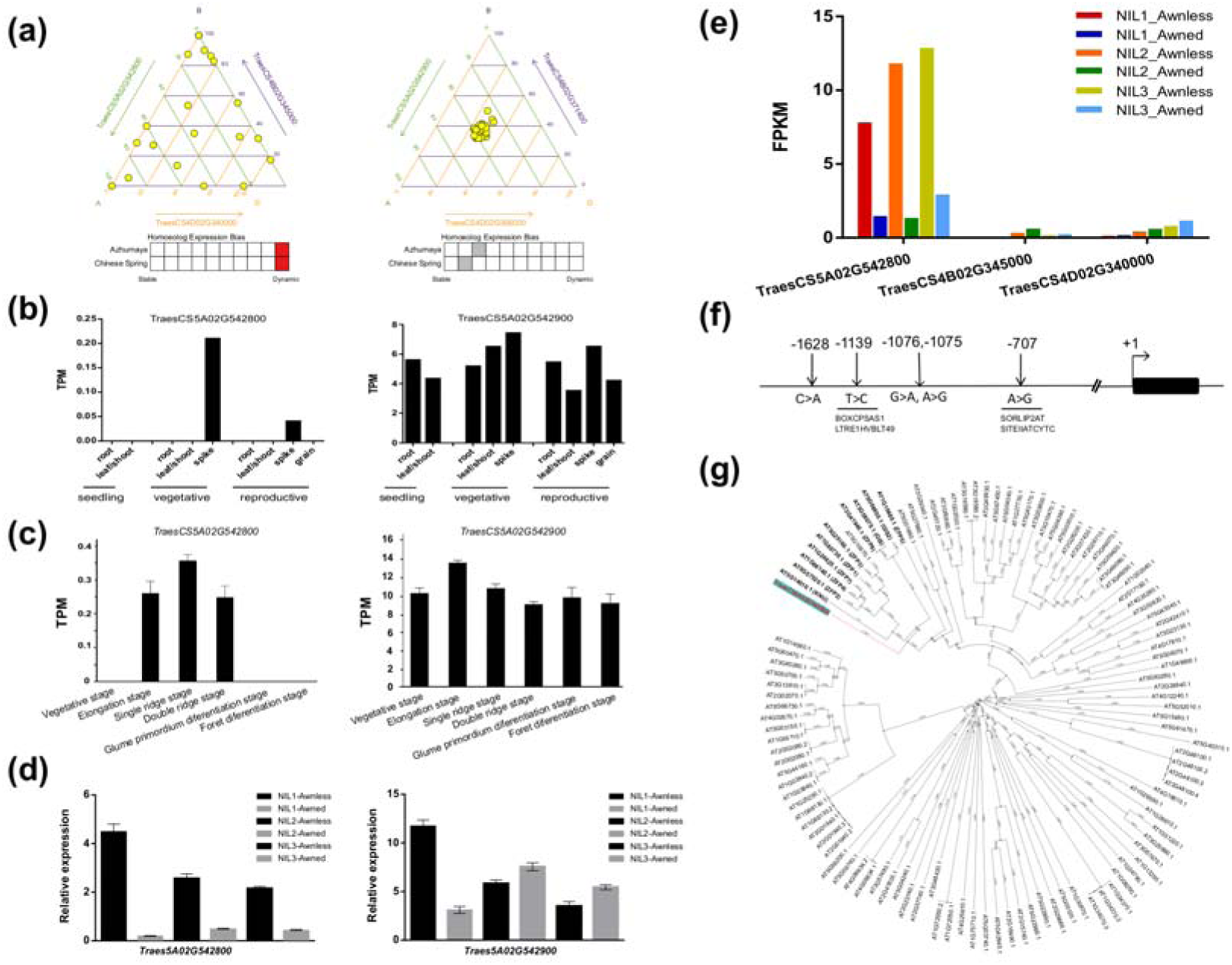
The expression patterns, sequence analysis, and phylogenetic analysis of candidate genes. (a) The time-course expression bias of homolog genes of *TraesCS5A02G542800* and *TraesCS5A02G542900* in Chinese Spring and Azhurnaya. Each yellow circle represents for the expression ratio of A, B, D homolog genes at a specific tissue and stage in CS or Azhurnaya. (b) The expression level of *TraesCS5A02G542800* and *TraesCS5A02G542900* in Chinese Spring at different tissue and stage. (c) The dynamic expression of *TraesCS5A02G542800* and *TraesCS5A02G542900* in KN9204 during the spike developmental process. (d) The relative expression levels of *TraesCS5A02G542800* and *TraesCS5A02G542900* in spikes of three pairs of NILs based on the internal control gene *Ta4045*. Error bars indicate SD. Each reaction was performed in three technical repeats. (e) The RNA-Seq data FPKM of homolog genes *TraesCS5A02G542800*, *TraesCS4B02G345000* and *TraesCS4D02G340000* in spikes of three pairs of NILs. (f) Consensus sequence variants identified between YMZ/S4185 and NK/ND3214 in the promoter region of *TraesCS5A02G542800*. (g) The neighbor-joining tree of *ALI-1* clustered with all C_2_H_2_ genes in *Arabidopsis thaliana*. The gene name marked in red (*TraesCS5A02G542800*) indicates *ALI-1*.

To determine the gene carrying the *B1* mutation, genomic DNA sequences of these two candidate genes, including exons, flanking intronic region, approximately 2-kb promoter region and 0.5-kb 3’-UTR region were sequenced in the parental lines of two mapping population, YMZ, S4185, NK and ND3214. Only five coincident SNPs in the promoter region of *TraesCS5A02G542800* were detected between the awned/awnless lines (**Figure 3f**). Among these five SNPs, the T>C mutation at -1139 bp could cause the loss of cis-elements BOXCPSAS1 and LTRE1HVBLT49 and the A>G mutation at -707 bp lost the cis-elements SORLIP2AT and SITEIIATCYTC (**Figure 3f**), and thus might be a causative factor for its significantly differential expression. No coincident polymorphisms in *TraesCS5A02G542900* were detected between YMZ/S4185 and NK/ND3214.

Meanwhile, using SMRT^®^ sequencing platform, *TraesCS5A02G542800* and *TraesCS5A02G542900* were sequenced to characterize sequence variations in 43 Chinese cultivars, 17 Chinese landraces, and 24 foreign accessions. No identical variants were detected in *TraesCS5A02G542900* and the coding region of *TraesCS5A02G542800*. However, a total of 31 variations (29 SNPs, a 25-bp deletion and a 1-bp insertion) were identified in the promoter region of *TraesCS5A02G542800* (**Table S9**). Among these variations, all the 10 accessions with *B1* allele (identified in the F_2_ population derived from the cross with long awn cultivar ShiAiYiHao) share the same haplotype for the 31 variations, but accessions with *b1* allele (27 long-awn accessions and 3 awnless accessions) have nine haplotypes, and the remaining 44 accessions with unknown allele have six haplotypes (**Table S9**). For the accessions with *b1* allele, they have a haplotype of CGAG at -1139, -1076, -1075 and -707 bp, expected for one Slovakia accession SV73. Of the 44 accessions with unknown allele, 39 accessions (88.64%) have the CGAG haplotype, and the left (11.36%) have the GAGA one. Noteworthy, this GAGA/CGAG haplotype coincides with the polymorphisms detected between the parental lines of our two mapping populations, YS and NN (**Figure 3f**). Thus, the GAGA/CGAG haplotype of *TraesCS5A02G542800* was highly consistent with its alleles of *B1* locus, which might be the causative factor for the differentiation of *B1*/*b1* allele.

According to the IWGSC RefSeq V1.1 annotation, *TraesCS5A02G542800* encodes a C_2_H_2_ zinc finger transcription factor protein. Protein sequences of *TraesCS5A02G542800* and all C_2_H_2_ genes in *Arabidopsis thaliana* were subjected to construct a neighbor-joining phylogenetic tree. *TraesCS5A02G542800* was grouped into the zf-C2H2_6 family (PF13912) together with cellular proliferation repressor *KNU*, trichome developmental regulators *ZFP5*, *ZFP7*, *ZFP8*, *GIS* and *GIS2*, and abscisic acid signaling negative regulators *ZFP1*, *ZFP2*, *ZFP3* and *ZFP4*, etc (**Figure 3g**).

In addition, *TraesCS5A02G542900* was excluded for its over-expressed transformants in an awned variety KN199 did not provide any awn-shortened or awnless performance. In summary, *TraesCS5A02G542800*, a predicted C_2_H_2_ zinc finger transcription factor, is a highly probable candidate gene for awn inhibitor *B1*, designated thereafter as *Awn Length inhibitor 1* (*ALI-1*).

### *ALI-1* negatively regulates awn elongation through restraining the cytokinin-mediated cell proliferation

To better understand the mechanism of *ALI-1* on awn development, RNA-Seq, paraffin sections, quantitative content determination of endogenous CKs and IAA were performed using three pairs of NILs. Transcriptome profiling of three pairs of NILs identified 1039, 2001, and 3387 differentially expressed genes (DEGs) between the awned and awnless individuals (**Figure 4a**). With this, a total of 478 overlapped DEGs were identified, including 244 up-regulated DEGs and 234 down-regulated ones in the awnless lines (**Figure 4b**). These overlapped DEGs were subjected to the analysis of GO enrichment and mapped to the reference KEGG pathways. The “nutrient reservoir activity”, “transporter activity”, “localization”, and “molecular transducer activity” were mostly enriched GO terms (**Figure 4c, Figure S4a**). For the KEGG, “phenylpropanoid biosynthesis”, “glutathione metabolism” and cytochrome P450 involved metabolism pathways were significantly enriched (**Figure S4b**). The up-regulated and down-regulated DEGs were separately subjected to GO enrichment analysis. “rRNA N−glycosylase activity”, “negative regulation of translation” and “defense response” were the most enriched GO terms in the up-regulated subgroup, and “circadian regulation of gene expression” and fatty−acyl−CoA metabolic process involved GO terms were also highly abundant (**Figure 4d**). In contrast, “transcription factor activity”, “regulation of cell size”, and auxin influx/efflux activity related GO terms were enriched in the down-regulated subgroup (**Figure 4e**). Notably, the up-regulated DEGs were most enriched in the cellular component of the Golgi membrane and chloroplast thylakoid membrane (**Figure 4f**), while the down-regulated ones located in plasmodesma and cell wall (**Figure 4g**). Similarly, some enriched GO terms in biological process (**Figure S4c,d**), molecular function (**Figure S4e,f**) and KEGG pathways (**Figure S4g,h**) were detected in the up- and down-regulated subgroup, respectively.

**Figure 4.**
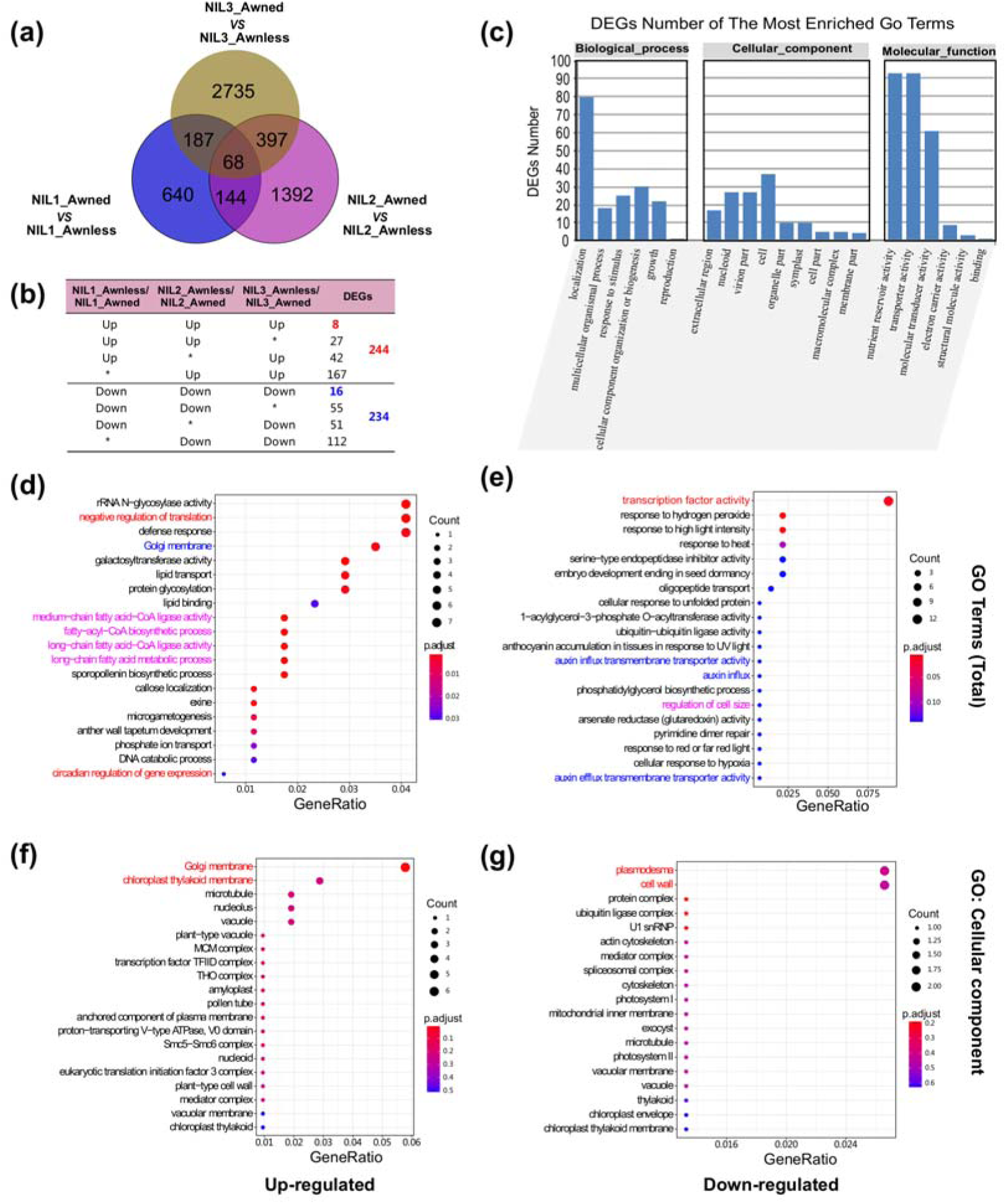
Transcriptome profiling of three pairs of NILs. (a) Differentially expressed genes identified from NIL1-Awnless *vs.* NIL1-Awned, NIL2-Awnless *vs* NIL2-Awned and NIL3-Awnless *vs* NIL3-Awned. (b) The number of genes with different expression pattern in three pairs of NILs. “Up”, “Down” and “*” indicates up-regulated genes, down-regulated genes and non-differentially-expressed genes in *ALI-1* lines, respectively. The numbers indicate gene numbers in each type. (c) The number of genes assigned to each GO terms in “biological process”, “cellular component” and “molecular function” of the 478 DEGs. (d,e) Dot plots of the top 20 GO terms of the up-regulated DEGs (d) and down-regulated DEGs (e) enriched in three pairs of NILs. GO terms were aligned with DEGs and considered to be significantly enriched with adjusting *P*-value < 0.05. The degree of enrichment was defined as the GeneRatio = N_GO Terms_/N_All DEGs_, in which N_GO Terms_ represents the number of DEGs in a specified GO term (“Count” in the GO plots) and N_All DEGs_ for the number of DEGs in all GO terms. (f,g) The dot plots of the top 20 GO terms enriched in the “cellular component” of the up-regulated DEGs (f) and down-regulated DEGs (g) enriched in three pairs of NILs.

Among the 478 overlapped DEGs, only eight genes including *ALI-1* (*TRIAE_CS42_5AL_TGACv1_374501_AA1201650*) were significantly up-regulated, and 16 genes were down-regulated in all three awnless NIL lines (**Figure 5a**). Since *ALI-1* is a transcriptional factor, a dual-luciferase reporter (DLR) assay system in Arabidopsis protoplasts was exploited to measure its transcriptional activation ability (Hao et al., 2010), using Pro*_35S_*:GAL4DBD:VP16 as a positive control (**Figure 5b**). Compared with the Pro*_35S_*:GAL4DBD negative control, Pro*_35S_*:GAL4DBD:ALI-1 decreases the luc activity by approximately seven folds (**Figure 5c**), providing a strongly transcriptional suppression activity of *ALI-1*. Therefore, the direct downstream target genes of *ALI-1* should exist in the 16 down-regulated gene set, including transcription factors *ZFP182* (*TRIAE_CS42_5BL_TGACv1_406541_AA1346800*) and *bHLH99* (*TRIAE_CS42_5AL_TGACv1_374213_AA1193840*), dual-specificity phosphatase *CDC25* (*TRIAE_CS42_5AL_TGACv1_377856_AA1249830*) and an auxin-responsive gene *IAA2* (*TRIAE_CS42_7DS_TGACv1_621699_AA2023540*) (**Figure 5a**). These genes were highly coincident with the “transcription factor activity”, “cell cycle arrest”, “cyclin-dependent protein serine/threonine kinase inhibitor activity” and “auxin influx/efflux activity” in the GO analysis of down-regulated genes.

**Figure 5.**
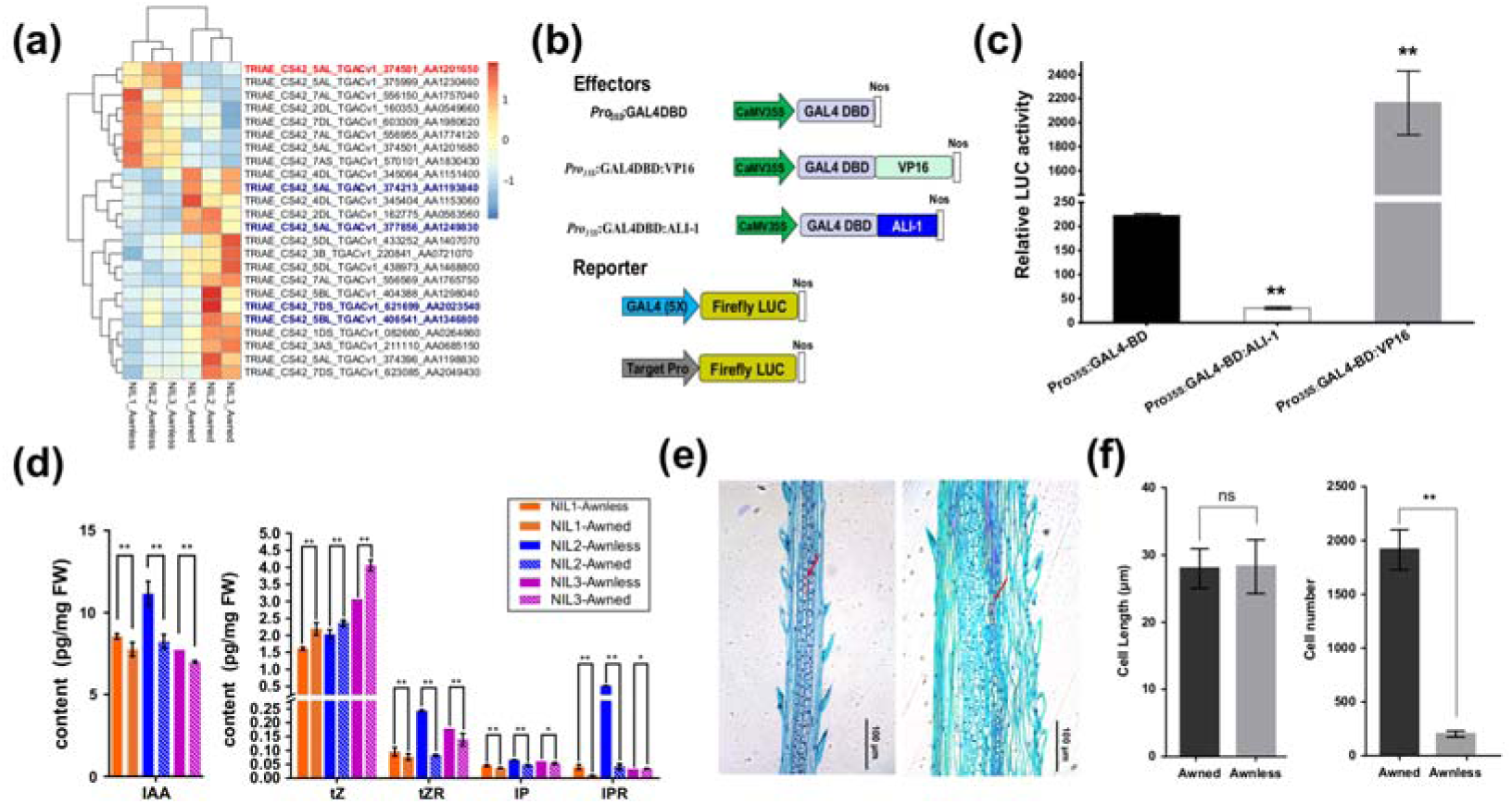
Transcriptional activation analysis, endogenous phytohormone quantitative analyses, and histological observation of *ALI-1*. (a) Heatmap of the 24 overlapped DEGs in three pairs of NILs. The expression level of each sample is print as deep blue representing the lowest value to deep red representing the highest value in the heat map. The gene marked in red indicates *ALI-1*, while genes marked in blue indicate possible direct downstream target genes. (b) Vectors used in the dual-luciferase reporter (DLR) assay system. (c) *ALI-1* strongly suppresses the luciferase activity of GAL4-LUC. The relative luciferase activity was measured, using the Pro*_35S_*:GAL4DBD and Pro*_35S_*:GAL4DBD:VP16 as a negative control and positive control, respectively. The error bars indicate SD from six independent measures of each analysis. (d) Endogenous IAA content and CKs contents in three pairs of NILs. The error bars indicate SD. One-way ANOVA test was used to determine the significance of the difference between awnless and awned lines. **, *P* < 0.01, *, *P* < 0.05. (e) The awn microscopic structure of awned (left) and awnless (right) plants. The red arrows indicate the cells used to the measurement of cell length and the irregular boxes depict the shape of cells. Bar = 100 µm. (f) The cell length and cell number in the awns of awnless and awned plants. The cell lengths of each sample were measured on three serial sections at the upper, middle, and bottom parts of awn. Cell number for the entire length of an awn was estimated based on the length of awns. Error bars show ± SD. One-way ANOVA test was used to determine the significance of the difference between awnless and awned lines. **, *P* < 0.01, ns, *P* > 0.05.

Endogenous IAA concentrations in the spike of three NILs were measured, as GO terms of auxin influx/efflux activity were significantly enriched in the down-regulated subgroup, and a significantly higher IAA content was observed in the *ALI-1* lines (awnless lines) (**Figure 5d**). For cytokinin, the concentration of *trans*-zeatin (*t*Z) was much higher than isoprenyl adenine (iP) in both the *ali-1* (awned lines) (47.32-folds) and *ALI-1* lines (32.41-folds) (**Figure 5d**), indicating that *t*Z was the main active cytokinin component in NILs. The concentrations of *t*Z in *ali-1* were higher than that in *ALI-1* (**Figure 5d**), which therefore might promote the division of awn primordium cells. On the contrary, nucleoside-type *trans*-zeatin nucleoside (*t*ZR) were lower in *ali-1* lines (**Figure 5d**), which might be owing to a dynamic transformation from *t*ZR to *t*Z. It is noteworthy that the type-A response regulator (K14492, ARR-A), a negative regulator in the cytokinin-mediated signal transduction, were significantly up-regulated in *ALI-1* lines (ko04075, https://www.kegg.jp/dbget-bin/www_bget?map04075). Hence, *t*Z was much less in *ALI-1* plants, and the cytokinin-mediated signal transduction was suppressed as the overexpression of negative regulator ARR-A.

To attribute the short awn of *ALI-1* to the cell size or cell number, longitudinal sections of awns in *ALI-1* (∼5 mm in length) and *ali-1* (∼50 mm in length) NILs were compared (**Figure 5e**). No significant differences in cell length detected between *ALI-1* (27.99 ± 0.56 μm) and *ali-1* (28.28 ± 0.85 μm) (**Figure 5f**). However, the longitudinal cell number in *ali-1* was nearly 10 times than that of *ALI-1* (**Figure 5f**), concluding that the cell proliferation was restrained in *ALI-1*, which might result from the reduced cytokinin content and/or suppressed cytokinin-mediated signal transduction.

### Pleiotropic effects of *ALI-1* on awn and grain development

The awn lengths in YS-F_2_, NN-F_2,_ and NILs were surveyed to evaluate the effect of *ALI-1* on awn performance (**Figure 2b**). The average AL of awned and awnless individuals in YS-F_2_ population was 49.63 mm and 4.51 mm, respectively, while it was 44.80 mm and 3.70 mm in Shi4185 and YMZ. Similar data were observed in the NN-F_2_ population, revealing the complete dominant characteristics of *ALI-1*. The value of AL reduction in three pairs of NILs was 54.69 mm, 62.30 mm and 54.11 mm, respectively, which was roughly equal to that in F_2_ populations but much higher than that of *qAL.5A.3_B1* in the open population of GWAS panel (reducing AL for 21.46 mm). As many loci were identified involving in the awn development, the effect of a single locus may be affected by the complicated additive-dominance-epistatic effects among those loci.

Other agronomic traits of these NILs were simultaneously measured in four environments (E1 and E2, normal condition of 2016–2017 and 2017–2018 growing seasons; E3 and E4, drought condition of 2016–2017, and 2017–2018 growing seasons). For most agronomic traits, there was no apparent difference between the *ALI-1* and *ali-1* individuals (**Figure S5a-j**), except for plant height under drought condition (**Figure S5k**) and kernel traits (**Figure 6a-h)**

**Figure 6.**
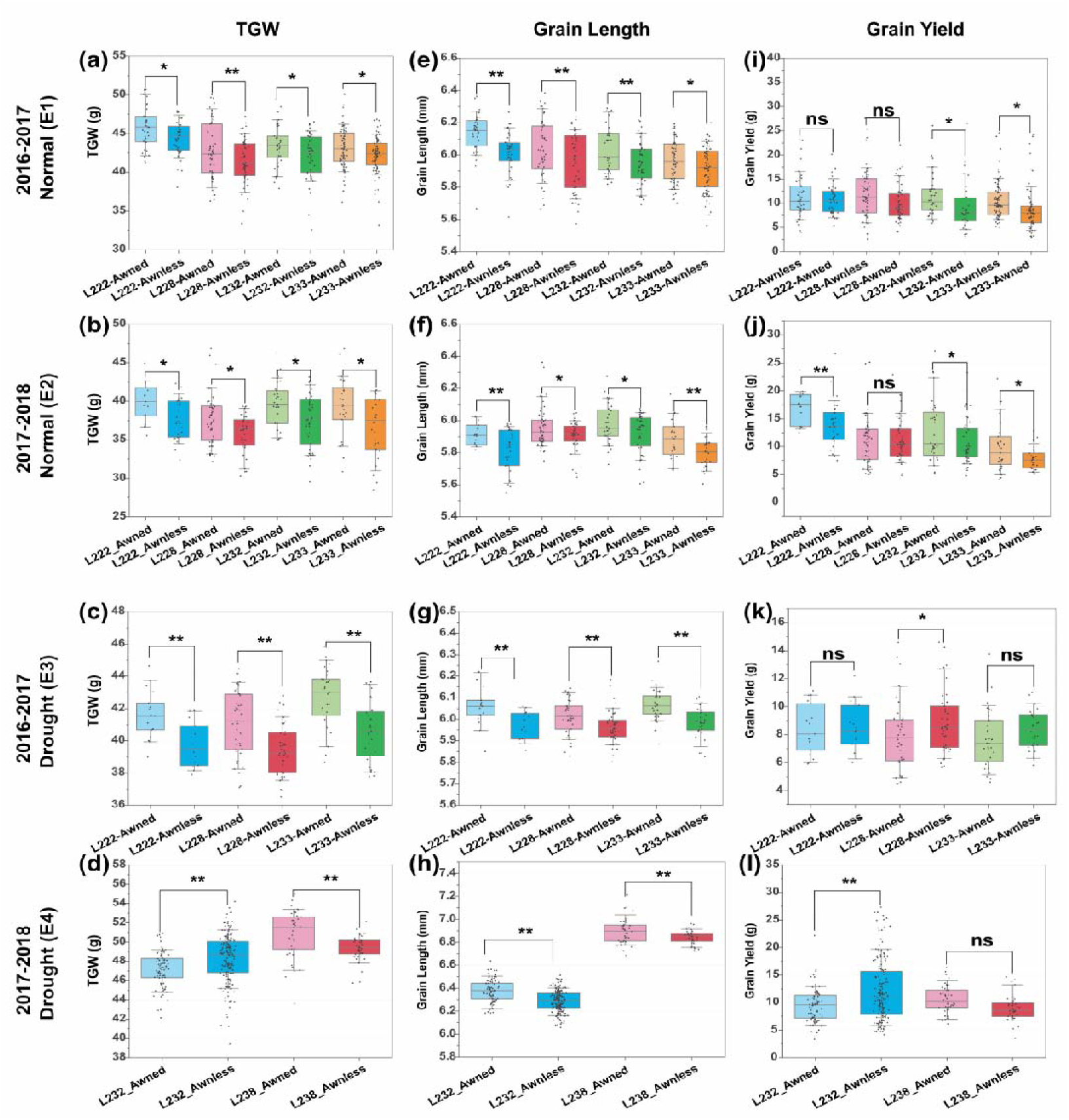
Effects of *ALI-1* on the yield-related traits. (a-l) The TGW (a-d), GL (e-h) and grain yield (i-l) performance of each line at environment E1–E4. The boxes cover the twenty-fifth to seventy-fifth percentiles with a middle line indicates median, the whiskers outside the box extend to the ±1.5 SD. TGW, GL and grain yield of individuals are displayed using gray dots. The significance of differences in TGW, GL and grain yield between awnless and awned lines were tested using one-way ANOVA. **, *P* < 0.01, *, *P* < 0.05, ns, *P* > 0.05.

*ALI-1* lines showed a significantly depressed thousand-grain weight (TGW) (**Figure 6a-d**). The TGW of four awnless NILs decreased ∼1.30 g under the normal condition, while that was up to 2.11 g under the drought condition, manifesting the contribution of *ALI-1* on wheat yield development, especially under the drought condition. Through the measurement of grain parameters, the reduction of TGW on the awnless NILs was attributed to the decrease of grain length (GL) (**Figure 6e-h**).

The contribution of awn on TGW was confirmed with our GWAS data. A significant correlation was observed between AL and TGW (Pearson’s *r^2^*=0.258) (**Figure 7a**). TGW associated SNPs were overlapping with the *qAL.5A.3_B1* locus, and the short awn haplotype-CCG had a TGW reduction of 5.31g (from 41.48g to 36.17g) as compared to the long awn haplotype-AAT. This reduction was consistent with the decrease of GL between two haplotypes (6.40 mm for haplotype-CCG and 6.24 mm for haplotype-AAT) (**Figure 7b**). Meanwhile, through re-assaying the available published data, the *qAL.5A.3_B1* locus was proven to contain the QTL for TGW and GL using recombinant-inbred line populations derived from the awnless×awned crosses (Li *et al.*, 2012; Wang *et al.*, 2011; Wu *et al.*, 2015).

**Figure 7.**
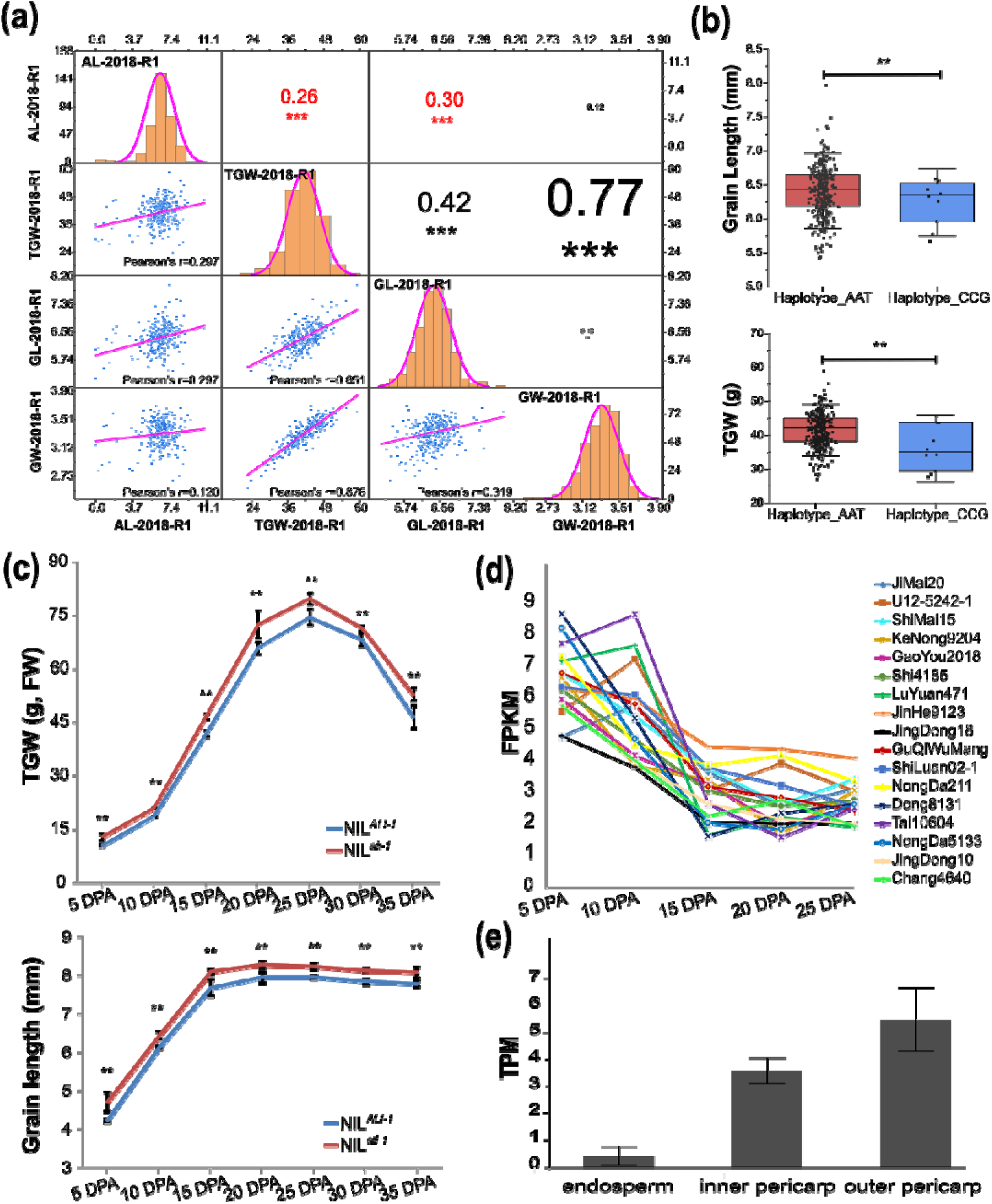
ALI-1 reduces TGW by suppressing grain length. (a) Distribution and correlation coefficients of AL (AL), grain length (GL), grain width (GW), and TGW of 2018-R1 in the GWAS panel. The frequency distribution of AL, GL, GW, and TGW was shown in the histogram at the diagonal cells. The X-Y scatter plot with the adjusted Pearson’s coefficients, and the corresponding Pearson’s coefficients between each trait were showed at the upper- and lower-triangle panel. ***, *P* < 0.001 in the multiple comparison significant test. (b) The grain length and TGW of haplotypes based on the genotype at *qAL.5A.3_B1* locus. The boxes cover the twenty-fifth to seventy-fifth percentiles with a middle line indicates median. The whiskers outside the box extend to the ±1.5 SD. GL and TGW of each individual are displayed using gray dots. The significance of differences in GL and TGW between different haplotypes was tested using one-way ANOVA. **, *P* < 0.01. (c) Comparison of the TGW (fresh weight) and grain length between the NIL*^ALI-1^* and NIL*^ali-1^* during grain development at 5, 10, 15, 20, 25, 30 and 35 DPA in 2018–2019 field trials. **, *P* < 0.01. The error bars indicate SD. (d) The dynamic expression of *bHLH99* during grain development in 17 Chinese cultivars. FPKM in the RNA-Seq data were used, two biological repetitions were carried out for each sample. (e) The expression level of *bHLH99* in different layers of the developing wheat grain at 12 DPA.

**Figure 8.**
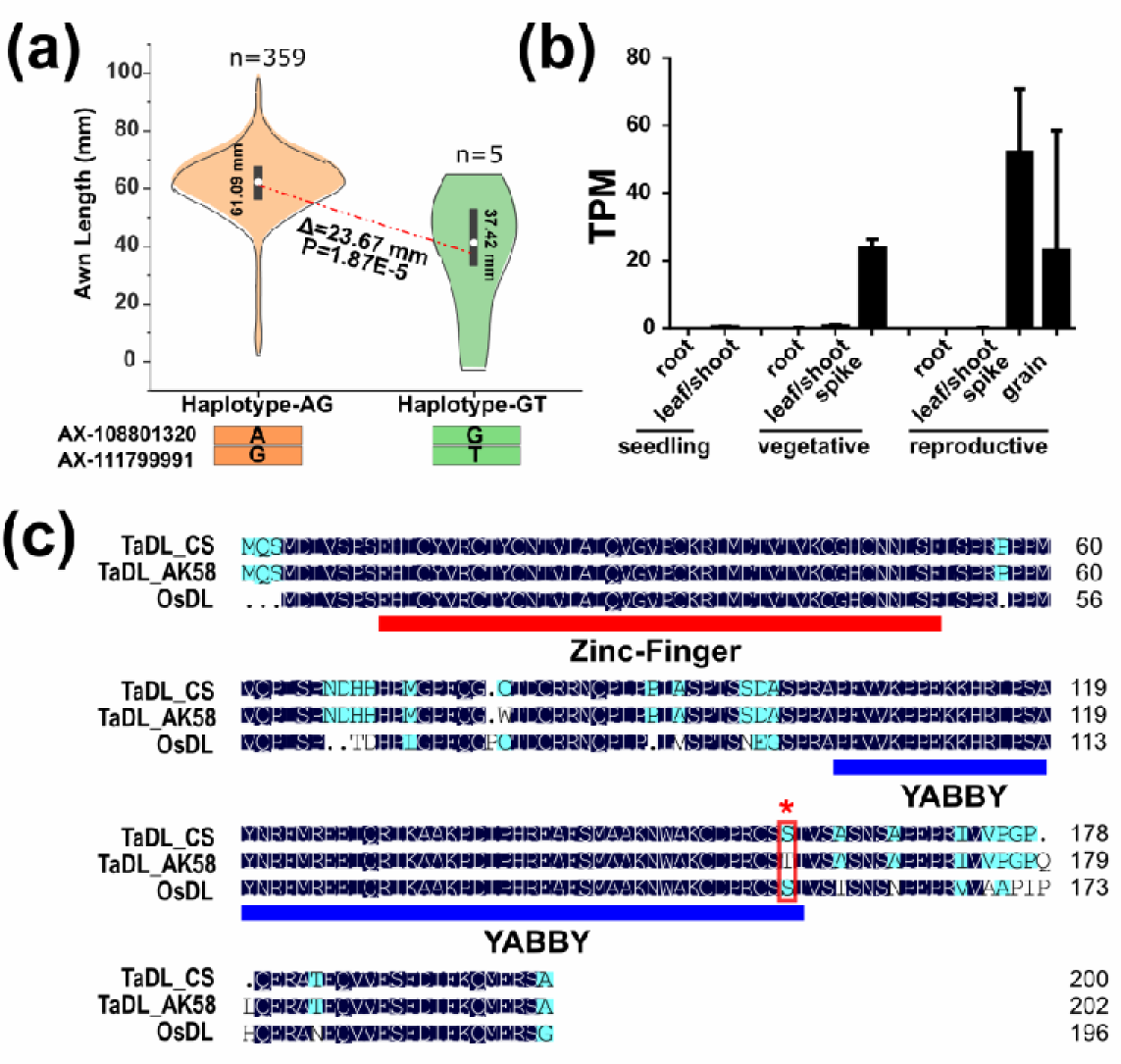
The haplotype analysis and candidate gene analysis of *Hd* locus. (a) The haplotype analysis of *Hd* locus. The violin plot with a standard box plot inside was used to compare the AL between two haplotypes, with mean values linked by a red dash line. One-way ANOVA test was used to determine the significance of difference, and the violin plot were plotted using software OriginPro, Version 2019. (b) The expression pattern of *TaDL* at different time/tissue in Chinese Spring. (c) Sequence alignment of the *DL* gene and its homolog *TaDL* in Chinese Spring (*TaDL_CS*) and AK58 (*TaDL_AK58*). The Zinc-finger domain and YABBY domain were annotated and the S>T amino acid substitution in AK58 indicated with an asterisk and a box.

Moreover, grain development time courses of NIL*^ALI-1^*/NIL*^ali-1^*were conducted and significant differences in the TGW was observed since the first investigate stage at 5 DPA (days post anthesis), suggesting the difference in final TGW between NIL*^ALI-1^*and NIL*^ali-1^* was caused by the grain development process rather than grain filling (**Figure 7c**). GL of NIL*^ALI-1^* and NIL*^ali-1^* rapidly increased during 5∼15 DPA and maintaining the status thereafter, and the difference of GL between NIL*^ALI-1^* and NIL*^ali-1^* was also detected throughout the time courses, indicating that *ALI-1* might affect the early grain development process (**Figure 7c**). Noteworthy, dynamic analysis of gene expression during grain development in 17 Chinese cultivars provides that *bHLH99* was highly expressed at 5 DPA and 10 DPA, but drastically reduced at 15 DPA and remained stable thereafter (**Figure 7d**). In addition, *bHLH99* was predominantly expressed in the pericarp, especially in the outer pericarp, of immature grain at 12 DPA (Pearce et al., 2015) (**Figure 7e**). Taken together, *ALI-1* might repress the expression of *bHLH99* in pericarp and consequently reduces the GL and TGW.

*ALI-1* was the first wheat awn controlling locus observed reducing GL and TGW, especially under drought condition. The contribution of awn to grain yield has been extensively researched, and the photosynthesis of awn was generally considered responsible for the improvement of grain weight (Grundbacher, 1963; Evans et al., 1972; Olugbemi et al., 1976; Li et al., 2006; Li et al., 2010). This work illustrates that *ali-1* removes sink limitation with larger grain size, and hence provides a reacquaint of the effect of wheat awn on grain production. Accordingly, regulating the expression of *ALI-1* and/or its downstream target genes would provide a strategy to achieve improved grain yield and address future extreme climate.

## Discussion

### Genome-wide identification of loci involved in wheat awn development

In this work, the GWAS provided 25 loci involved in AL on 14 chromosomes, among which six was overlapped with known QTL in wheat or wheat homologs of awn controlling genes in rice and barley. *Lks2* in barley was the first cloned gene for awn length in the grass family. The short-awn *lks2* allele is present in limited accessions and was a natural variation that occurred after barley domestication (Yuo et al., 2012). A stable SAL on chromosome 7A detected in all environments was overlapped with wheat homolog of *Lks2*, explaining 9.52% phenotypic variation of BLUP value and reducing AL of 8.97 mm, which was medium compared to other loci and accorded with its incomplete baldness in barley (**Figure 1c**, **Table S5)**. Unlike its low frequency of *lks2* allele in barley, short awn haplotype AAG included 91 accessions (25.00%) in GWAS panel, indicating that the long-awn allele *Lks2* has not been artificially selected during the domestication. The *An-1*, regulating long awn formation in *O. rufipogon*, was a major target for artificial selection in rice (Luo et al., 2013). However, the elite haplotype of its wheat homolog comprises 59 accessions (16.21%) and with a weak effect in wheat (reduces 3.85 mm of AL). *DL* affects the formation of rice awn and *OsETT2* enhances its elongation, while *SHL2* acts on *OsETT2* transcripts to inhibit the awn length (Toriba and Hirano, 2014). The short awn allele of *OsETT2* and *SHL2* homologs were detected in a limited proportion of accessions (20 and 26 accessions, respectively), suggesting an artificial selection during the breeding history. Two SNPs at the genome region of *DL* homolog was significantly associated with the AL, which were insufficient to form a SNP cluster and not observed as a SAL, and was overlapped with the important wheat awn inhibitor *Hd* (Yoshioka et al., 2017). The haplotypes AG and GT comprising 359 accessions and 5 accessions, respectively, with a 23.67 mm difference in AL (61.09 ± 11.99 mm *vs* 37.42 ± 20.29 mm) (**Figure 7a**). The wheat homolog of *DL* is predominantly expressed in the spike of Chinese Spring (**Figure 7b**), and an S>T amino acid substitution at the conserved YABBY domain was found in the long-awn variety AK58 (**Figure 7c**), suggesting that *DL* might be the candidate gene conferring to the awn inhibition of *Hd* locus. Hence, homologs of several genes controlling the awn development in rice and barley affect the AL in GWAS panel, and these genes exhibit to be functionally conserved and might experience parallel evolution/domestication across different species.

Chinese Spring deletion line 5AL-10 was reported slightly bearded while 5AL-17 was awnless (Sourdille et al., 2002), but no QTL was detected on chromosome 5AL using a doubled-haploid line population derived from the cross of Courtot and Chinese Spring, which confused researchers for a long time (Sourdille et al., 2003; Yoshioka et al., 2017). The deletion line 5AL-10 and 5AL-17 lack the telomeric region of the long arm of chromosome 5A, with breakpoints located between Xgwm156-Xgwm617 and Xcfa2163-Xcfa2155, respectively (Sourdille et al., 2004; Yoshioka et al., 2017). In this work, we identified a new locus *qAL.5A.2* (547.59–548.25 Mb) located between the breakpoints of 5AL-10 and 5AL-17 (Xgwm156-Xcfa2155, 450.16-632.60 Mb). Thus, the awn-suppressing *qAL.5A.2* allele might locate in the 5AL-17, and the deletion of this locus in the 5AL-10 relived its inhibition on awn development. The q*AL.5A.2* reduced the AL by 23.02 mm in the GWAS panel (**Figure 1c**, **Table S5**), which was equivalent to the AL of line 5AL-10. However, that the *qAL.5A.2* was not detected in the population of Courtot and Chinese Spring might due to the lack of enough markers in the genetic linkage map surrounding *qAL.5A.2* locus, or other loci interacted/complemented with the *qAL.5A.2* in Courtot to result in an awn performance.

### *ALI-1* represses cytokinin-mediated cell proliferation in awn

*ALI-1* encodes a C_2_H_2_ zinc finger transcription factor protein, and the phylogenic analysis grouped it with cellular proliferation repressor *KNU* and trichome developmental regulators *ZFP5*, *ZFP7*, *ZFP8*, *GIS*, and *GIS2* (**Figure 3g**). *KNU* is a transcriptional repressor of cellular proliferation in *Arabidopsis* (Payne et al., 2004). C_2_H_2_ zinc finger proteins integrate hormonal signals to control trichome cell differentiation in *Arabidopsis*, and *GIS2*, *GIS3*, *ZFP5*, *ZFP6*, and *ZFP8* were reported to regulate trichome initiation through GA and cytokinin signaling (Gan et al., 2007; Sun et al., 2015).

*ALI-1* seems to have a similar role in controlling the awn elongation, suppressing the cytokinin signaling and cell proliferation. In the NILs, concentrations of *t*Z in *ali-1* were higher than that of *ALI-1* lines (**Figure 5d**), and even the cytokinin signal transduction was suppressed because of the overexpression of negative regulator ARR-A. The stimulatory effect of cytokinin was achieved through cytokinin-mediated cell cycling arrest of plant tissues, and the plant homolog of *CDC25* was considered as an early target for cytokinin action (John, 1998; Lipavská, H. et al., 2010). We screened the ∼2000 bp promoter regions of the 16 down-regulated DEGs to search for the binding sequence A[AG/CT]CNAC of C_2_H_2_ zinc finger proteins (Sun et al., 2015). One, Two, and three perfect matches were detected in the promoter region of *CDC25*, *bHLH99*, and *IAA2*, respectively. *CDC25* was the only gene that expression changes highly resembling that of *ALI-1*, NIL-2 > NIL-1 > NIL-3, with an average 28.1-folds down-regulation in awnless individuals. Thus, the absence of CDC25 accumulation in *ALI-1* lines might further aggravate the inadequate cytokinin signal on promoting cell division. Longitudinal sections of awns showed that the numbers of cells were markedly decreased in the awns of *ALI-1* lines (**Figure 5e,f**). Besides, a prominent enrichment of GO terms with cell cycle, plasmodesma, and cell wall were obtained in the down-regulated DEGs. Sequence analysis provides that SNPs in the promoter region lead to the absence of cis-elements BOXCPSAS1, LTRE1HVBLT49, SORLIP2AT and SITEIIATCYTC in *ALI-1* (**Figure 3f**), which is involved in the regulation of gene expression in meristematic tissues and/or proliferating cells (Hudson, 2003; Welchen, 2006).

Taken together, we speculate that SNPs in the promoter of awnless individuals result in the up-regulation of *ALI-1* and the consequent trace expression of *CDC25*, and this reduces the cytokinin content and simultaneously restrains the signal transduction of cytokinin, which leads to a stagnation of cell proliferation and reduction of cell number. As a consequence, the elongation of awn in *ALI-1* was inhibited and presented as very short awn phenotype. Due to a cascading effects of transcription factor on the downstream genes, *ALI-1* exhibits an exceeding inhibition on awn elongation that plant carries this allele (even in heterozygous state) to be awnless without the presence of *B2* or *Hd*. However, it’s still unclear whether and how *ALI-1* directly regulates cytokinin concentrations.

### *ALI-1* pleiotropically regulates awn and grain development

The long spiculate awn with barbs severely hinders manual harvesting and storage; however, as a potential photosynthetic organ, awn could significantly increase the grain weight, especially under drought condition (Evans et al., 1972; Li et al., 2010). Except enhancing photosynthesis source, *ali-1* also acts to remove sink limitation, providing a larger grain size, and might manipulate the carbon source-sink balance. *NSG*, the *ALI-1* homolog in rice, involved in the regulation of glume length (Wang et al., 2013), which was the key physiological factor limiting grain size in rice. In our NILs and GWAS population, an appreciable effect of *ALI-1* on GL and TGW was observed, which is consistent with previously reported QTL (Li *et al.*, 2012; Wang *et al.*, 2011; Wu *et al.*, 2015). Moreover, analysis of grain growth process in the GWAS panel detected associated SNPs within the *ALI-1* region at 5 DPA and 10 DPA (unpublished), indicating that *ALI-1* involves the grain formation at the lag phase (Bennett *et al.*, 1975). *bHLH99* showed significant sequence similarities with *OsRHL1*, *An-1*, and *PIL* genes, and it was predominantly expressed at the early grain development process, especially in the pericarp (**Figure 7d,e**). *An-1* prolongs cell division in the lemma, resulting in an increased cell number and grain length (Luo et al., 2013). *PGL1* mediates the grain elongation and increases grain weight by controlling cell elongation in lemma and palea (Heang and Sassa, 2012). Besides, *OsPIL1*/*OsPIL13* regulates internode elongation and plant height via cell wall-related genes in response to drought stress in rice (Todaka et al., 2012). Under drought condition, plant height was decreased and a notable increase in GL was obtained in *ali-1*. In addition, plasmodesma and cell wall were the most enriched cellular components GO terms in the down-regulated DEGs (**Figure 4g**). Taken together, *ALI-1* might negatively regulate the expression of *bHLH99* in the developing grains, resulting in a reduction of GL and TGW in awnless lines. Thus, silencing *ALI-1* and regulating its downstream target genes would theoretically increase the awn length and accordingly broaden the photosynthesis source and kernel sink simultaneously, which would provide an alternative strategy to improve wheat yield potential. Nevertheless, a better understanding of its mechanism in regulating the grain elongation is pre-requisite before its practical application in the wheat breeding program.

## Supplementary data

Figure S1 Distribution and correlation coefficients of AL for the six environments in the GWAS panel.

Figure S2 Manhattan plots and quantile-quantile plots of AL for the six environments and the BULP value.

Figure S3 Genome regions showing strong association signals and the LD plot around the *qAL.5A.3_B1* locus.

Figure S4 Top enriched GO terms and KEGG pathways in the up- and down-regulated genes.

Figure S5 Phenotypic performances of spike length, spikelet number per spike, plant height, grain width, spike number, and grain number per spike in the NILs.

Table S1 Accessions used in the Genome-Wide-Association Study and their awn performance.

Table S2 Phenotype variation of awn length in the six environments.

Table S3 Analysis of variance of awn length in the GWAS panel.

Table S4 Significantly associated loci of wheat awn length identified in the GWAS.

Table S5 Descriptive statistics and ANOVA of awn length between haplotypes.

Table S6 Annotation of genes in the *qAL.5A.2* LD block.

Table S7 SNPs associated with wheat awn length around the *qAL.5A.3_B1* locus.

Table S8 The χ^2^ test of awn segregation of the four populations used in the mapping of *B1* locus.

Table S9 Haplotype analysis of *ALI-1* in 84 accessions.

Table S10 Primers used in this study.

## Acknowledgements

This work was supported financially by National Key Research and Development Program of China (2016|YFD0101004, 2016YFD0101802) and National Natural Science Foundation of China (31571643).

## Author’s contributions

DL and AZ conceived and supervised the study; DW, DL, KY, DJ, LS, JC, WW, WY, JS, XL and PX conducted the research and analyzed the data; DW and DJ collected phenotypic data; DW, KY, DL and KZ conducted the GWAS; DW, KY, DJ and WW participated in the fine mapping; DW collected samples for RNA-Seq and quantitative content determination of endogenous CKs and IAA. DW and KY contributed to analyses of transcriptomic data; PX, JC conducted the quantitative content determination of endogenous CKs and IAA, and JC, DW analyzed the data; DW and DJ carried out the paraffin section; DJ and DW contributed to the determination of transcriptional activation ability. DW, DL, and AZ prepared the manuscript. All authors discussed the results and commented on the manuscript.

