## Supplemental figure S1-S5 for "*ALI-1*, candidate gene of *B1* locus, is associated with awn length and grain weight in common wheat"

### Supplemental data

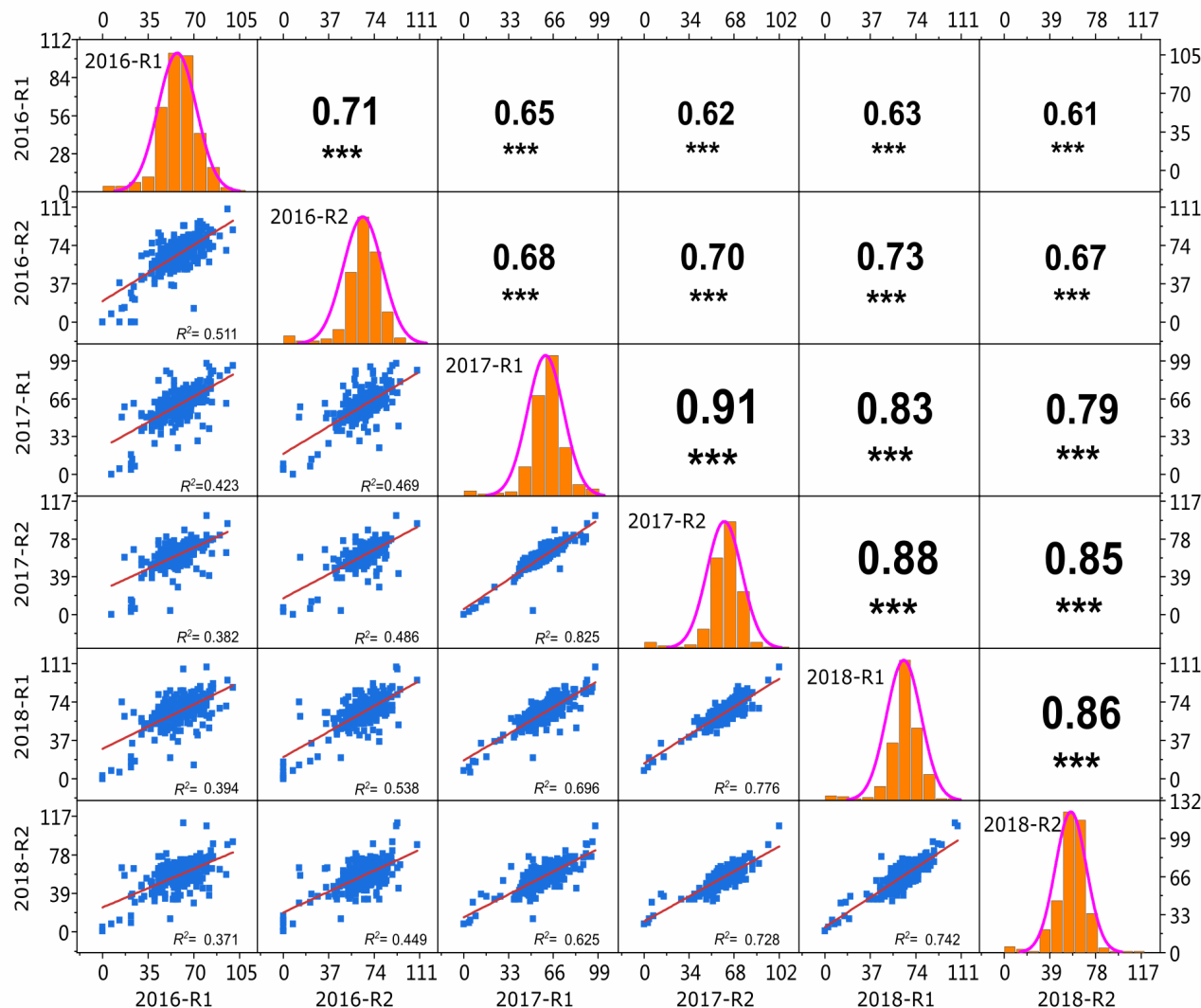
**Figure S1 Distribution and correlation coefficients of AL for the six environments in the GWAS panel.** The frequency distribution of AL at each environment was shown in the histogram at the diagonal cells. The X-Y scatter plot showing the correlation between environments at the lower-triangle panel, while the corresponding Pearson’s coefficients between each trait were showed at upper-triangle panel. ***, *P* < 0.001.

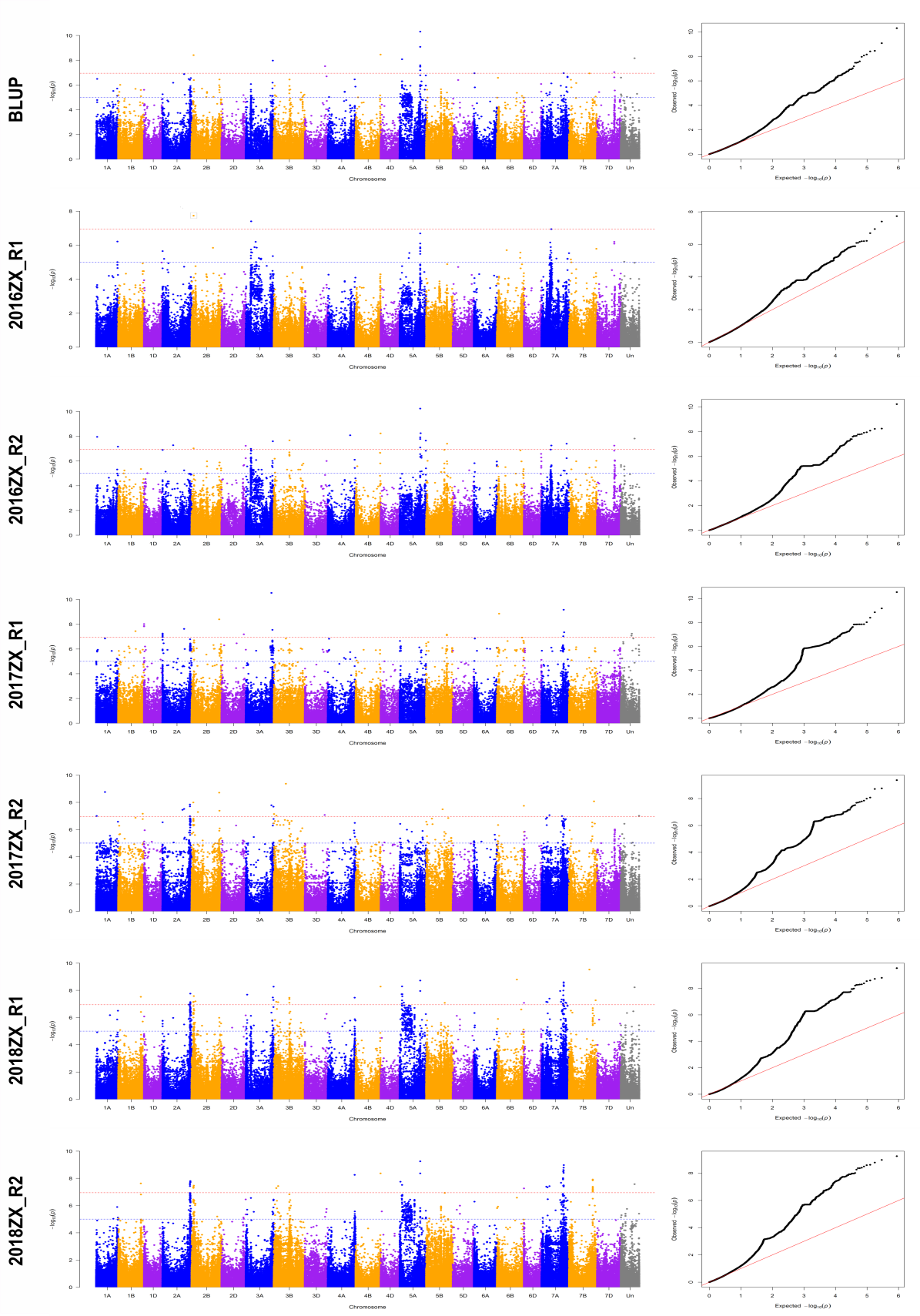

**Figure S2 Manhattan plots and quantile-quantile plots of AL for the six environments and the BULP value.** Manhattan plots for AL identifies SAL across the 21 chromosomes using the mixed linear model (MLM). The -log_10_(*P*) values from a genome-wide scan are plotted against positions on 21 chromosomes. Blue and red horizontal dashed lines indicate the genome-wide significance threshold of *P*=10^−5^ and *P*=10^−7^, respectively. The solid red line in quantile-quantile plot indicates the expected values.

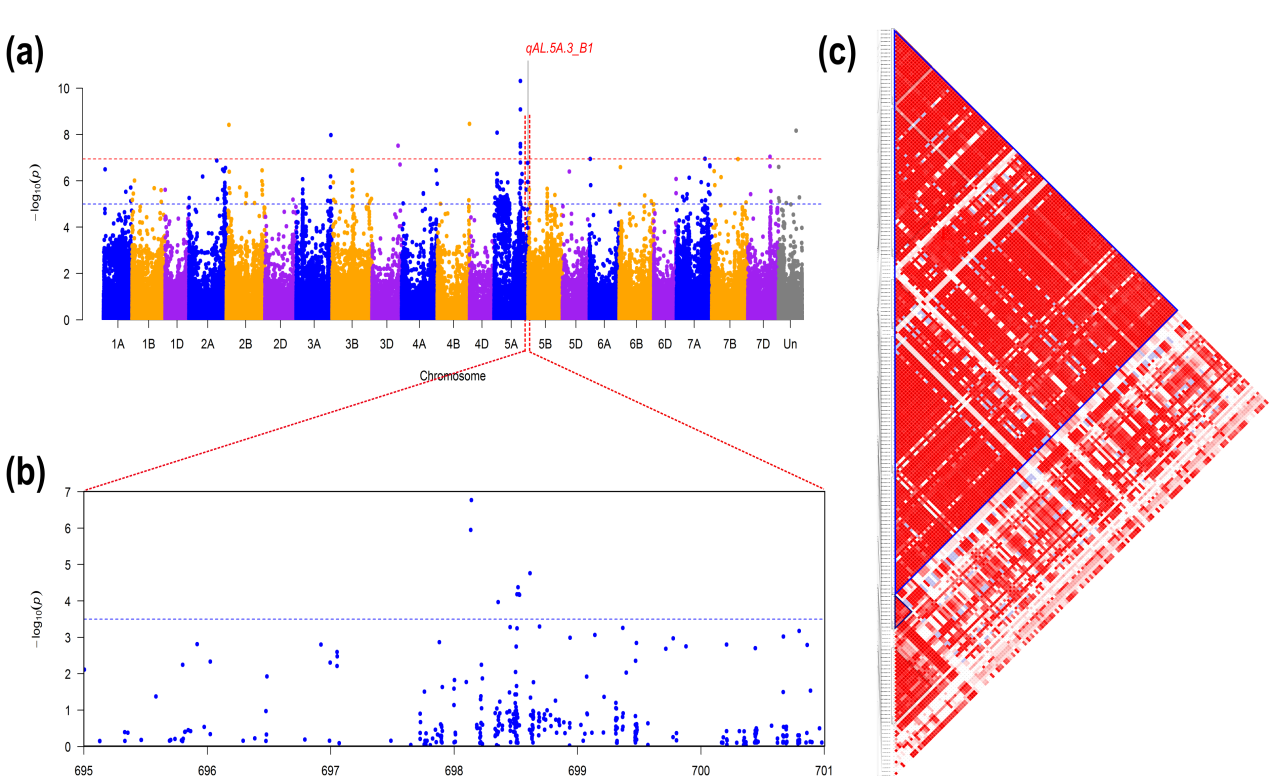

**Figure S3 Genome regions showing strong association signals and the LD plot around the *qAL.5A.3_B1* locus.** (a) Manhattan plots for BULP value of AL using the mixed linear model (MLM) (b) SNPs in 6-Mb region flanking the peak SNP (with the lowest *P* value) of *qAL.5A.3_B1* locus were displayed. The blue horizontal dashed lines indicate the manual-set genome-wide significance threshold of *P*=0.0005. (c) LD plot of the 2 Mb region across the peak SNP. The formed LD blocks were indicated using a blue triangular frame.

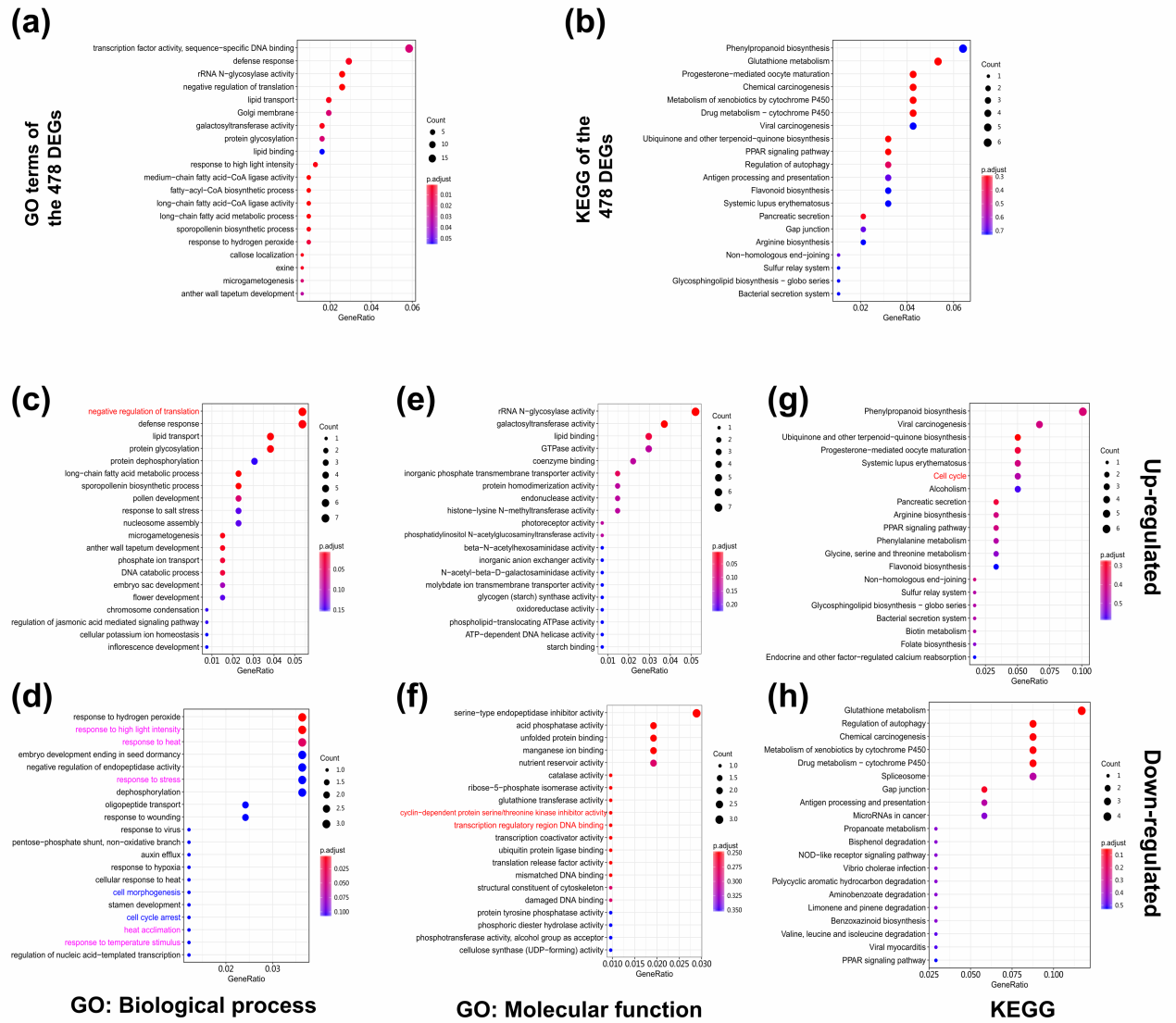

**Figure S4 Top enriched GO terms and KEGG pathways in the up-regulated genes and down-regulated genes.** (a) The top 20 enriched GO terms of the 478 DEGs. (b) The top 20 enriched KEGG pathways of the 478 DEGs. (c,d) The top 20 enriched GO terms of type “biological process” in the up-regulated genes and down-regulated genes. (e,f) The top 20 enriched GO terms of type “molecular function” in the up-regulated genes and down-regulated genes. (g,h) The top 20 enriched KEGG pathways in the up-regulated genes and down-regulated genes.

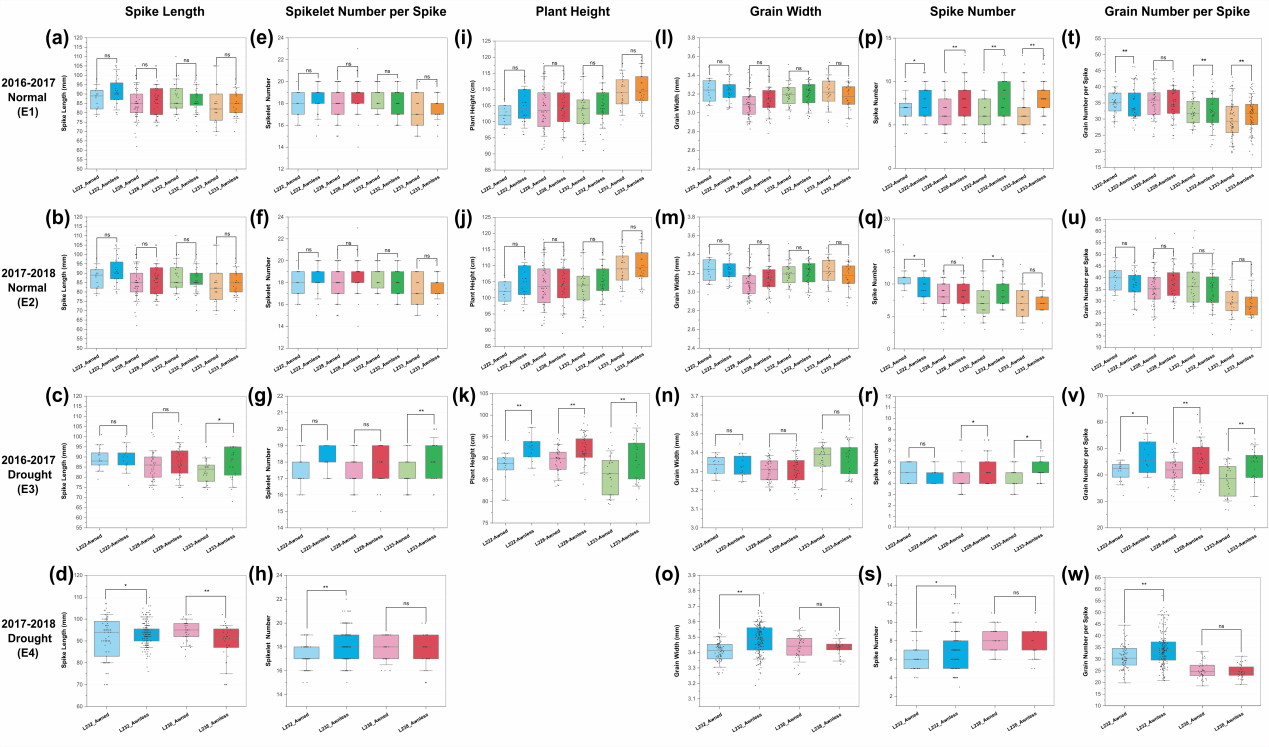

**Figure S5 Phenotypic performances of spike length, spikelet number per spike, plant height, grain width, spike number and grain number per spike in the NILs.** The performances of spike length (a-d), spikelet number per spike (e-h), plant height (i-k), grain width (l-o), spike number (p-s) and grain number per spike (t-w) of each line in the NILs at environment E1–E4 were displayed use the box plot. The boxes cover the twenty-fifth to seventy-fifth percentiles with a middle line indicates median, the whiskers outside the box extend to the ±1.5 SD. Data from each individuals are displayed using gray dots. The significance of differences between awnless and awned lines were tested using one-way ANOVA. **, *P* < 0.01, *, *P* < 0.05, ns, *P* > 0.05. The Tukey's original box plot were plotted using software OriginPro, Version 2019.
